# RNA sequencing and lipidomic analysis of alveolar macrophages from normal and CD44 deficient mice

**DOI:** 10.1101/845644

**Authors:** Yifei Dong, Arif A. Arif, Jian Guo, Zongyi Ha, Sally S. M. Lee-Sayer, Grace F. T. Poon, Manisha Dosanjh, Calvin D. Roskelley, Tao Huan, Pauline Johnson

## Abstract

Alveolar macrophages (AMs) are CD44 expressing cells that reside in the alveolar space where they maintain lung homeostasis by serving critical roles in immunosurveillance and lipid surfactant catabolism. AMs lacking CD44 are unable to bind the glycosaminoglycan, hyaluronan, which compromises their survival and leads to reduced numbers of AMs in the lung. Using RNA sequencing, lipidomics and multiparameter flow cytometry, we demonstrate that CD44^-/-^ mice have impaired AM lipid homeostasis and increased surfactant lipids in the lung. CD44^-/-^ AMs had increased expression of CD36, a lipid scavenger receptor, as well as increased intracellular lipid droplets, giving them a foamy appearance. RNA sequencing revealed the differential expression of genes associated with lipid efflux and metabolism in CD44^-/-^ AMs. Lipidomic analysis showed increased lipids in both the supernatant and cell pellet extracted from the bronchoalveolar lavage of CD44^-/-^ mice. Phosphatidylcholine species, cholesterol, oxidized phospholipids and levels of reactive oxygen species (ROS) were increased in CD44^-/-^ AMs. Oxidized phospholipids were more cytotoxic to CD44^-/-^ AMs and induced greater lung inflammation in CD44^-/-^ mice. Reconstitution of CD44^+/+^ mice with CD44^-/-^ bone marrow as well as adoptive transfer of CD44^-/-^ AMs into CD44^+/+^ mice showed that lipid accumulation in CD44^-/-^ AMs occurred irrespective of the lung environment, suggesting a cell intrinsic defect. Administration of colony stimulating factor 2 (CSF-2), a critical factor in AM development and maintenance, increased AM numbers in CD44^-/-^ mice and decreased phosphatidylcholine levels in the bronchoalveolar lavage, but was unable to decrease intracellular lipid accumulation in CD44^-/-^ AMs. Peroxisome proliferator-activated receptor gamma (PPARγ), downstream of CSF-2 signaling and a regulator of lipid metabolism, was reduced in the nucleus of CD44^-/-^ AMs, and PPARγ inhibition in normal AMs increased their lipid droplets. Thus, CD44 deficiency causes defects in AMs that lead to abnormal lipid accumulation and oxidation, which exacerbates oxidized lipid-induced lung inflammation. Collectively, these findings implicate CD44 as a regulator of lung homeostasis and inflammation.

## INTRODUCTION

Alveolar macrophages (AMs) are important for lung homeostasis, playing key roles in immunosurveillance and lipid surfactant catabolism (1). They are fetal monocyte-derived tissue resident macrophages (2) in the alveolar space that require colony stimulating factor 2 (CSF-2 also known as GM-CSF) (2, 3), peroxisome proliferator-activated receptor gamma (PPARγ) (4), and transforming growth factor beta (TGFβ) (5) for their development, maturation and survival. The deficiency of any of these factors leads to the accumulation of immature macrophages which are unable to catabolize surfactant lipids, and this leads to the buildup of lipids in the immature AMs which take on a foamy appearance. Consequently, lung surfactant lipids build up in the alveolar space and this can lead to pulmonary alveolar proteinosis (PAP) (6, 7). While primary PAP is attributed to defective CSF-2 signaling, the causes of secondary PAP are largely unknown (7). CSF-2 induces the expression of PPARγ in fetal lung monocytes, which is critical for the generation of the AM gene signature (4). PPARγ is also important for lipid catabolism (8-10) in mature AMs (4). TGFβ upregulates PPARγ expression and conditional deletion of the TGFβ receptor in macrophages leads to the accumulation of immature, lipid-laden foamy macrophages, showing a key role for TGFβ signaling in AM development (5). Autocrine TGFβ, produced by mature AMs, also contributes to their self-maintenance and survival (5).

AMs have an important role in the clearance of pulmonary surfactant produced by alveolar epithelial cells which is typically made up of 80% polar lipids (primarily phosphatidylcholine (PC)), 10% neutral lipids including cholesterol, and 10% proteins (1, 7, 11). AM numbers are reduced during lung damage and inflammation (12-14) and this leads to the accumulation of surfactant lipids in the alveolar space where they become susceptible to oxidation in the oxygen-rich environment of the lung, especially during inflammation when reactive oxygen species (ROS) are generated by inflammatory cells (15). Oxidized lipids are cytotoxic, and if they build up in AMs, lung function is compromised (16). For example, bleomycin-induced lung injury leads to inflammation, ROS production, phospholipid oxidation and the generation of foamy macrophages, which all contribute to lung dysfunction (16). In contrast, mice lacking a NADPH-oxidase subunit that cannot generate ROS were protected from bleomycin-induced pulmonary damage and fibrosis (17). In humans, oxidized lipid cytotoxicity can lead to the life-threatening acute respiratory distress syndrome (18) and lung phospholipid peroxidation is associated with asthma and chronic obstructive pulmonary disease progression (19). Furthermore, oxidized phosphatidylcholine (OxPC) is found in AMs from patients diagnosed with idiopathic interstitial pneumonia (20). Thus, oxidized lipids are an important component of the pathophysiology of lung disease, and the removal of excess surfactant lipids and oxidized lipids from the alveolar space by AMs is essential to maintain vital lung function and homeostasis.

Unlike most other macrophages at steady state, mature murine AMs express high levels of CD11c, Siglec F, and constitutively bind hyaluronan (HA) via its receptor, CD44 (21, 22). HA, a component of the extracellular matrix, is implicated in both lung homeostasis and inflammation (23). At homeostasis, CD44-dependent HA binding by AMs results in the formation of a pericellular HA coat that promotes their survival (12). AMs from CD44^-/-^ mice lack the HA coat, leading to a decrease in their viability and CD44^-/-^ mice have approximately half the normal number of AMs (12). In humans, CD44 expression is lower on AMs isolated from patients with diffuse panbronchiolitis (24) and chronic obstructive pulmonary disease (COPD) (25), and in mice, the depletion of AMs exaggerates pulmonary inflammatory responses (26, 27). Furthermore, CD44^-/-^ mice treated with bleomycin are unable to resolve lung inflammation, resulting in a buildup of HA and apoptotic neutrophils (28, 29). Here, we investigated the impact of CD44 in lung homeostasis and inflammation by studying AMs from CD44^-/-^mice.

## MATERIALS AND METHODS

### Mice

C57BL/6J (CD45.2^+^), B6.SJL-Ptprc^a^Pepc^b^/BoyJ (CD45.1^+^), C57BL/6J x BoyJ heterozygotes (CD45.1^+^, CD45.2^+^), and CD44^-/-^ mice were housed and bred in specific pathogen free facilities, as previously described (12). Experiments used 6 to 14 week-old, age and sex matched mice and were conducted with protocols approved by the University Animal Care Committee in accordance with the Canadian Council of Animal Care guidelines for ethical animal research.

### Reagents

Fluorescein-conjugated HA (FL-HA) was prepared as previously described (12). *Streptomyces hyaluronlyticus* hyaluronidase (HA’se) and biotinylated HA-binding protein (HABP) were from Millipore, and streptavidin was from Thermo Fisher Scientific or eBioscience. Recombinant mouse CSF-2 (carrier free) was from BioLegend. L-α-phosphatidylcholine (PC) was from Sigma-Aldrich. 1-palmitoyl-2-(5’-oxo-valeroyl)-sn-glycero-3-phosphocholine (POVPC) was from Avanti Polar Lipids. Fatty acid free (FAF) BSA was from Roche Diagnostics. BODIPY 493/503, N-(7-Nitrobenz-2-Oxa-1,3-Diazol-4-yl)-1,2-Dihexadecanoyl-sn-Glycero-3-Phosphoethanolamine (NBD-PE), and 2’,7’-dichlorodihydrofluorescein diacetate (H2DCFDA) were from Thermo Fisher Scientific. The PPARγ antagonist, T0070907, was from Tocris Bioscience. Fluorescent or biotinylated labeled CD11c (N418), CD11b (M1/70), Ly6C (HK1.4), CD14 (SA2-8), CD45.1 (A20), CD45.2 (104), CD200R (OX110), MHC II (M5/114.15.2), Siglec F (1RNM44N) and SIRPα (P84) were from Thermo Fisher Scientific; CD36 (HM36) and CD206 (C068C2) were from Biolegend; CD44 (IM7) was from Ablab; Ly6G (1A8), Siglec F (E50-2440) and mouse IgG1κ isotype were from BD Biosciences; CD116 (698423) and MerTK (polyclonal BAF591) were from R&D; PPARγ (81B8) was from Cell Signaling Technologies, and anti-OxPC (E06) was from Avanti Polar Lipids.

### *In vivo* experiments

Bone marrow (BM) reconstitution into irradiated recipients was essentially as described in (12). POVPC (200 µg) in 50 µl PBS, or CSF-2 (2 µg daily) was given to mice by intratracheal instillation (i.t.) by laryngoscopic manipulation, and the bronchoalveolar lavage (BAL) analyzed 3 or 7 days later, respectively. For adoptive transfer experiments, 2-3 × 10^5^ CD44^+/+^ or CD44^-/-^ AMs were transferred by i.t. into CD45.1^+^ CD44^+/+^ or CD45.2^+^ CD44^-/-^ mice, and BAL was analyzed on day 7.

### Isolation of bronchoalveolar lavage (BAL) cells for RNAseq, flow cytometry or *in vitro* analysis

BAL was isolated from CD44^+/+^ and CD44^-/-^ mice using 4 × 700 µl PBS, 2 mM EDTA and the BAL cells (>95% AMs from naïve mice) were treated by RBC lysis buffer (0.84% NH4Cl in 10 mM Tris pH 7.2) for 5 min, centrifuged and either subjected to RNA extraction using RNeasy Mini Kit (Qiagen) or counted manually using the hemocytometer and trypan blue, and prepared for flow cytometry as previously described (12). Alternatively, AMs were washed once with PBS and incubated with 2.5 µM BODIPY 493/503 in PBS, 5 µM H2DCFDA in PBS, or 10 µM NBD-PE in RPMI for 30 min at 37°C prior to flow cytometry. Or, AMs were incubated with 1% EtOH (vehicle solvent), 50 µM PC, or 50 µM POVPC in PBS for 1 h at 37 °C, then analyzed with Annexin V and DAPI by flow cytometry (12). HABP was used to detect cell surface HA by flow cytometry (12).

### *In vitro* AM culture

AMs were cultured with 20 ng/ml of recombinant CSF-2 in RPMI containing 1% BSA, in the presence or absence of 500 µM PC or 1 µM T0070907 in a 96-well non-tissue culture treated plate for 48 h, harvested using versene, and the cells analyzed by flow cytometry. For culture with PC, 1% FAF BSA was used.

### Intracellular labeling of AMs

AMs were isolated by BAL with PBS 2 mM EDTA and fixed with 2% paraformaldehyde immediately for 10 min at room temperature, washed twice with PBS, labeled with the E06 or PPARγ antibody intracellularly using the Intracellular Fixation & Permeabilization Buffer Set (Thermo Fisher Scientific).

### Analysis of BAL fluid

BAL isolated using PBS 2 mM EDTA was centrifuged and the supernatant and cell pellet collected. Hyaluronan Quantikine ELISA Kit (R&D Systems) was used to determine the HA concentration in the BAL supernatant. Total protein was determined using the Pierce BCA Protein Assay Kit (Thermo Fisher Scientific). Total PC concentration was determined using the Phospholipase C assay kit (Wako Diagnostics).

### Gas chromatography analysis for cholesterol

Lipids were extracted twice from 1 ml BAL supernatant by adding 6 ml of 2:1 v/v chloroform: methanol, with 1% acetic acid. Organic solvents were evaporated under a stream of nitrogen gas. Lipid extracts were resuspended in pyridine and then an equal part of N,O-bis(trimethylsilyl)trifluoroacetamide with trimethylchlorosilane. Samples were spiked with 10 µM cholestane as an internal standard and analyzed by gas-chromatography mass spectrometry (GC-MS).

### Flow cytometric analysis

Flow cytometry was analyzed using FlowJo VX (Treestar). Flow cytometry plots shown were first gated by size, singlets, and live/dead stain. AMs were then gated as CD11c^+^ and Siglec F^+^ (Supplementary Figure S1A-B). Fold difference was calculated by dividing each CD44^+/+^ and CD44^-/-^ biological replicate by the mean of CD44^+/+^ samples in each experiment.

### Confocal microscopy

AMs labeled with BODIPY or E06 and DAPI were mounted onto Superfrost^+^ microscope slides (Thermo Fisher Scientific). Whole cell image stacks were acquired using a Leica Sp8 microscope. Maximum stack projections were analyzed using ImageJ 1.51 (NIH). BODIPY stained lipid droplets were identified using a trainable segmentation function in ImageJ, followed by particle analysis, and droplet masks were applied onto the raw images to measure number, pixel intensity, and droplet size. E06 labeled droplets were also counted. PPARγ and DAPI labeled AMs were acquired using a Leica Sp5. Total nuclear PPARγ was measured within the DAPI^+^ nuclear area and quantified as mean pixel intensity (MPI). Immunofluorescence of 12 µm lung tissue cryosections was measured as previously described (12) and analyzed using ImageJ for HABP mean pixel intensity (MPI), where HA-rich bronchioles were excluded from the analysis.

### RNAseq analysis

RNA from three independent CD44+/+ and three CD44^-/-^ samples were pooled from the BAL of 4, 2 or 2 female mice respectively.

#### Illumina RNA sequencing

RNA was sequenced by the UBC Biomedical Research Center. RNA quality, *18S* and *28S* ribosomal RNA with RIN = 9.6, was determined by Agilent 2100 Bioanalyzer following the standard protocol for NEBNext Ultra II Stranded mRNA (New England Biolabs). Sequencing was performed on the Illumina NextSeq 500 with paired end 42bp × 42bp reads. De-multiplexed read sequences were then aligned to the *Mus musculus* (mm10) reference sequence using Spliced Transcripts Alignment to a Reference, STAR (https://www.ncbi.nlm.nih.gov/pubmed/23104886), aligners. The sequence data is available at GEO database, accession number: GSE138445

#### Cufflinks and Cuffdiff

Assembly and differential expression were estimated using Cufflinks (http://cole-trapnell-lab.github.io/cufflinks/) through Cufflinks Assembly & DE version 2.1.0 which included Cuffdiff analysis (30). Gene differences with a q value (p value adjusted for false discovery rate) less than 0.05 were considered significant.

#### Metascape analysis

Metascape analysis was performed on the genes annotated as significantly different from the Cuffdiff analysis (31). Gene IDs were entered and analyzed as *M. musculus*. Significantly different GO pathways were obtained as an output from Metascape.

#### R Scripts and Heatmaps

R language was used for bioinformatic analysis of RNA sequencing results using RStudio Version 1.1.442. Gene ID was converted to Entrez ID using Annotables: Ensembl 90 conversion chart (32). Heatmaps were created using R library, “pheatmap()”. The input for heatmaps were a list of differentially expressed genes in specific GO pathways and their corresponding fragments per kilobase of transcript per million mapped reads (FPKMs). FPKM values were obtained from Illumina RNA-Sequencing.

#### String analysis

All 200 significant genes (q < 0.05) were input into the String Consortium (33). String Consortium maps the interactions between genes based on text mining, experiments, databases, co-expression, neighbourhood, gene fusion, and co-occurrence.

### Mass spectrometry, lipidomics and analysis

#### BAL cell pellet and supernatant sample preparation

The BAL from 8 CD44^+/+^ and 8 CD44^-/-^ female mice were spun down to generate a supernatant and cell pellet for each sample. Lipids were extracted from the BAL pellets, CD44^+/+^ (WT1-8) and CD44^-/-^ (KO1-8) with 400 µl ice cold extraction solvent (methanol:acetonitrile (ACN):water=2:1:1, v:v:v) followed by three freeze-thaw cycles. Proteins were precipitated at −20°C for 2 h, then 900 µl methyl tert-butyl ether (MTBE) was added and vortexed for 90 min on an open-air microplate shaker. 170 µl H2O was then added into the solution followed by centrifugation (17,530 g, 4°C, 1 min) to accelerate the phase separation. CD44^+/+^ (WT1-8) and CD44^-/-^ (KO1-8) BAL supernatant samples (3 ml each) were thawed on ice and 3 ml MTBE added and vortexed for 90 min on an open-air microplate shaker. In both cases, the upper organic phase was carefully transported to a 1.5 ml Eppendorf vial and concentrated by Speed-vac at 4°C. The concentrated pellets were subsequently reconstituted with ACN:isopropyl alcohol(IPA)=1:1, v:v for LC-MS analysis. One method control (MC) was prepared using the same procedure but with no sample in the vial. One quality control (QC) was prepared by pooling 10 µl from each of the 16 reconstituted solutions (8 WT+8 KO). QC was injected between every 4 sample injections to ensure the instrument was stable and the sample injection volume was optimized.

#### Liquid chromatography mass spectrometry (LC-MS) analysis

The LC-MS analysis was performed on Bruker Impact II™ UHR-QqTOF (Ultra-High Resolution Qq-Time-Of-Flight) mass spectrometer coupled with a Agilent 1290 Infinity™ II LC system. LC separation was performed on a Waters ACQUITY UPLC BEH C18 Column (130Å, 1.7 µm, 1.0 mm × 100 mm). 2 µl sample solutions of each replicate, MC and QC were injected in a random order. 2 µl Sodium formate was injected within each run for internal calibration. The mobile phase A was 60% ACN 40% H2O (2 mM ammonium acetate (NH4Ac) in positive mode, 5 mM NH4Ac in negative mode); mobile phase B was 90% IPA 10% ACN (2 mM NH4Ac in positive mode, 5 mM NH4Ac in negative mode). The chromatographic gradient was run at a flow rate of 0.100 ml/min as follows: 0-8 min: linear gradient from 95% to 60% A; 8-14 min: linear gradient from 60% to 30% A; 14-20 min: linear gradient from 30% to 5% A; 20-23 min: hold at 5% A; 23-24 min: linear gradient from 5% to 95% A; 24-33 min: hold at 95% A. The mass spectrometer was operated in Auto MS/MS mode. The ionization source capillary voltage was set to 4.5 kV in positive scanning mode and −3.5 kV in negative scanning mode. The nebulizer gas pressure was set to 1.0 bar. The dry gas temperature was set to 220°C. The collision energy for MS/MS was set to 7 eV. The data acquisition was performed in a range of 50-1200 m/z at a frequency of 8 Hz.

#### Data interpretation

Bruker Data Analysis software was used to calibrate the spectra using sodium formate as the internal reference. Then the raw data files with format of ‘.d’ were converted to format of ‘.abf’ using AbfConverter. The converted files were then uploaded onto MS-DIAL for lipid feature extraction and alignment. Lipids identification was carried out using LipidBlast available in MS-DIAL (34, 35). The parameters were set as follows: MS1 tolerance was 0.01 Da; MS2 tolerance was 0.05 Da; retention time tolerance was 0.5 min. The alignment result was exported in a ‘csv.’ file. Fold change (FC, calculated as CD44^-/-^/CD44^+/+^) and p values between CD44^+/+^ and CD44^-/-^ groups were calculated using a two tails distribution, homoscedastic *t* test in Microsoft Excel. The lipids with FC>1.5 or FC<0.67, and p value<0.05 were selected as being significantly different between the CD44^+/+^ and CD44^-/-^ groups.

### Statistics

Statistical analysis used is described in the figure legends. The majority of graphs were generated using GraphPad Prism 6 and the data shown are the average ± standard deviation. Significance was defined as * p < 0.05, ** p < 0.01, *** p < 0.001.

## RESULTS

### CD44^-/-^ AMs have increased cellular granularity and cell surface receptor expression

CD44^-/-^ mice have approximately 50% less AMs compared to normal C57Bl/6J mice, as previously reported (12). These CD44^-/-^ AMs express the characteristic AM markers: CD11c and Siglec F, but are unable to bind hyaluronan, an extracellular matrix glycosaminoglycan that promotes their survival (12). Here, flow cytometric analysis showed that CD44^-/-^ AMs, gated as in Supplemental Figure 1A-B, were unable to bind fluorescein labeled HA (FL-HA), had a greater granularity (SSC) and a slightly higher autofluorescence than CD44^+/+^ AMs (Figure 1A-B). In addition, there was a slight increase in the mean fluorescence intensity (MFI) of several cell surface molecules normally expressed on resident AMs (CD11c, CD206, CD200R, Sirpα, MerTK, PD-L1) but Siglec F levels remained unchanged (Figure 1C-D and Supplemental Figure S1C-D). Adjusting for increased autofluorescence did not account for these changes. There was also increased expression of molecules not normally expressed on mature AMs (CD36, MHCII, CD11b, and PD-1) (Figure 1C-D and supplemental Figure S1C-D). CD36 is a scavenger receptor for fatty acids and oxidized lipids (36, 37), raising the possibility that the increased SSC and autofluorescence may represent increased lipid content. To determine if cell surface HA bound on normal AMs physically affected the binding of antibodies and thus contributed to the phenotypic differences observed by flow cytometry, CD44^+/+^ and CD44^-/-^ AMs were treated with hyaluronidase (HA’se). While the binding of HABP on CD44^+/+^ AMs was reduced to CD44^-/-^ levels after HA’se treatment, there was no change in the MFI of cell surface receptors (Supplemental Figure S1E-F). These differences suggest CD44 deficiency causes changes in AMs.

**Figure 1.**
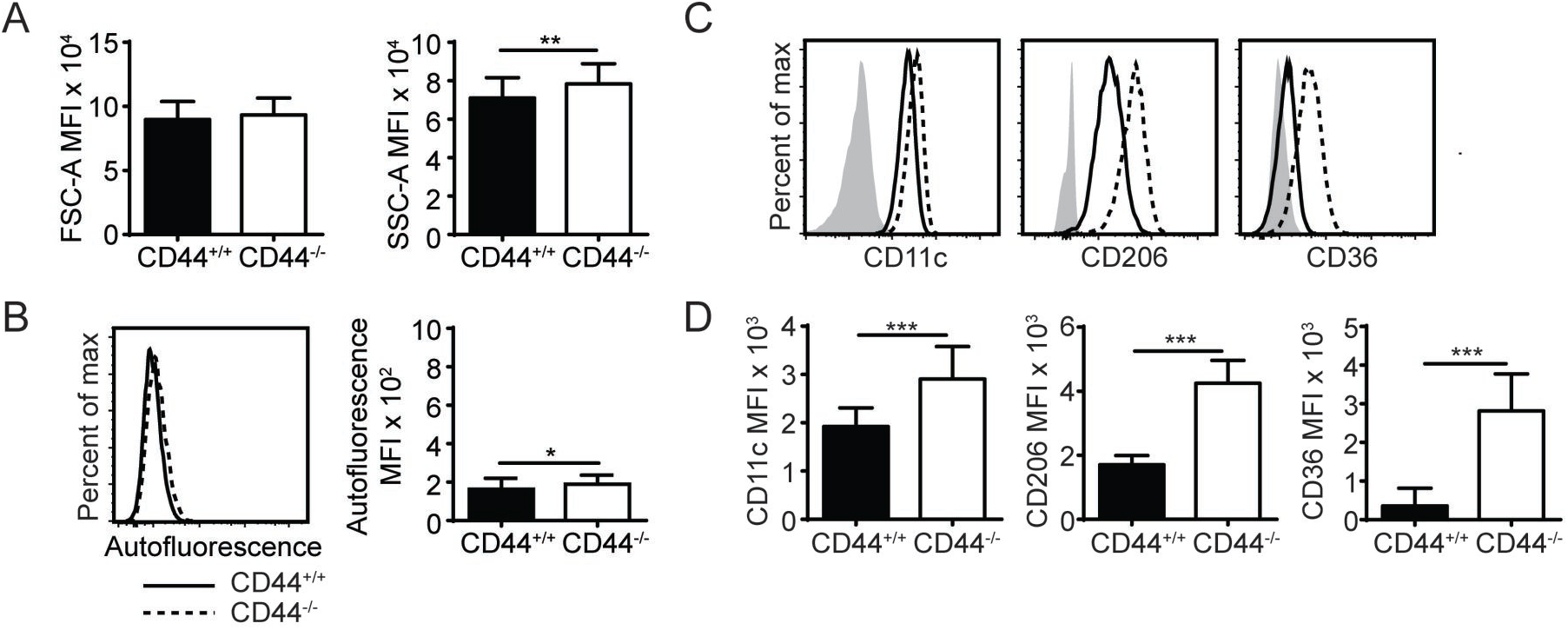
CD44^-/-^ mice have abnormal AMs. **(A)** and **(B)** Bar graphs and representative flow cytometry histograms comparing the FSC-A (cell size), SSC-A (cell granularity) and autofluorescence of CD44^+/+^ and CD44^-/-^ AMs from BAL of naïve mice. **(C)** and **(D)** Representative flow cytometry histograms and graphs comparing the MFI of cell surface phenotype of CD44^+/+^ and CD44^-/-^ AMs, where the background MFI of autofluorescence from unlabeled CD44^+/+^ or CD44^-/-^ cells was subtracted from the values shown. Data show an average of two experiments ± SD, each with three to five CD44^+/+^ and CD44^-/-^ mice, except for in **(A)** which was the average of BAL AMs from forty-four CD44^+/+^ and CD44^-/-^ mice over twelve experiments. Significance indicated as * p< 0.05, ** p< 0.01, *** p < 0.001, non-paired Student’s t-test.

### CD44^+/+^ and CD44^-/-^ AMs have different transcriptional profiles at steady-state

To understand how the loss of CD44 could be impacting AMs, we performed RNAseq analysis on CD44^+/+^ and CD44^-/-^ AMs. Approximately 200 genes with a q value of 0.05 or less were differentially expressed (Figure 2A and Supplemental Table 1). The RNAseq data showed several transcriptional changes in CD44^-/-^ AMs that could affect lipid homeostasis in the CD44^-/-^ AMs. Gene ontology analysis was performed to determine the pathways most affected by the loss of CD44 and this identified the greatest changes in the ‘sterol metabolic process’, followed by differences in ‘the response to external stimulus’, ‘protein phosphorylation pathways’, ‘regulation of homeostatic process’ and ‘regulation of sterol metabolic process’ (Figure 2B-C). String analysis of the differentially expressed genes led to the identification of a gene cluster associated with sterol/cholesterol synthesis, and a broader cluster of genes involved in cell signaling, adhesion and migration (Supplemental Figure S2). Dissection of the sterol pathways led to a significant number of up and down-regulated genes associated with sterol/cholesterol synthesis and trafficking. Upregulated genes included those involved in lipid/cholesterol transport (*ApoE, ApoC1, ABCA1*), triacylglycerol synthesis (*DGAT2)*, lipid catabolism (*LipN*), signaling (*Src, Igf1, Igf2r*), and immune function/antigen presentation (*CD74, MHCII*). Down regulated genes included phospholipase (*PLC)A2γ7, PLCβ1, Plin1, Srebf2, LDLR*, all involved in lipid homeostasis or its regulation. The transcription factor, Srebf2, is one of the master regulators of cholesterol homeostasis. In keeping with this, several genes in sterol/cholesterol synthesis were also downregulated (Figure 2D). The down regulation of genes involved in cholesterol synthesis and LDL uptake, and the upregulation of genes involved in sequestering lipids/cholesterol (*ApoE*) and trafficking them out of the cell (*ABCA1*), raised the possibility that the CD44^-/-^ AMs were responding to increased lipids/cholesterol.

**Figure 2.**
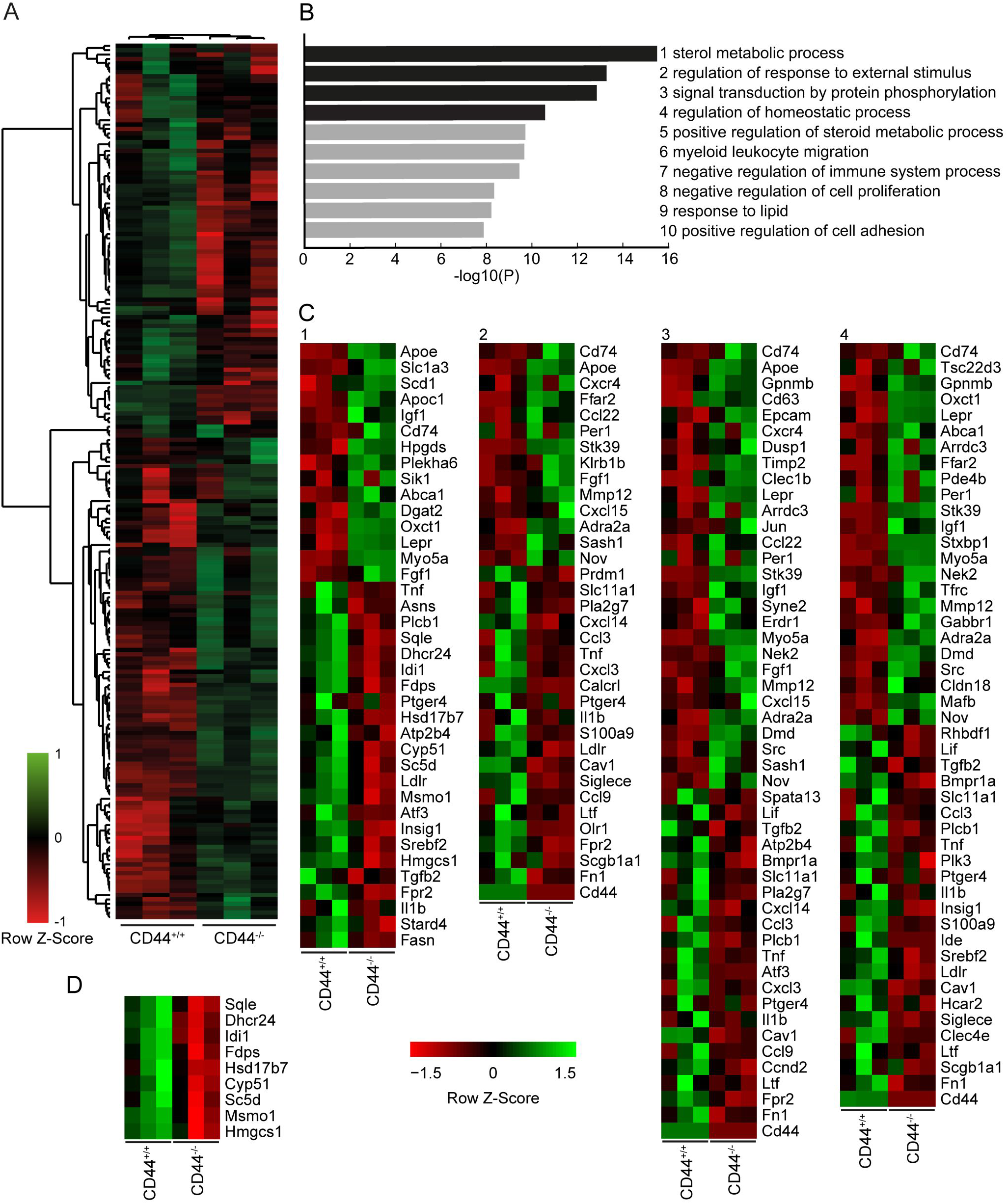
RNA sequencing analysis comparing CD44^+/+^ and CD44^-/-^ AMs. **(A)** Cuffdiff RNA sequencing analysis showing 200 significant differentially expressed genes (q-value < 0.05) between CD44^+/+^ and CD44^-/-^ AMs. 87 genes were decreased, and 113 genes were increased in CD44^-/-^ AMs. **(B)** Top 10 most significant Gene Ontology pathways associated with the significant differentially expressed genes analyzed by Metascape. **(C)** Heatmap of normalized FPKM of differentially expressed genes in the top 4 pathways of Metascape. **(D)** Heatmap of normalized FPKM of differentially expressed genes associated with the Cholesterol Biosynthesis Process, R-MMU-191273. Data are from 3 independent samples from a total of 8 mice for both CD44^+/+^ and CD44^-/-^ AMs.

### Lipidomic analysis reveals altered lipid levels in the BAL supernatant and cell pellet of CD44^-/-^ mice

To investigate if the transcriptional changes in CD44^-/-^ AMs affected lung lipid surfactant homeostasis in CD44^-/-^ mice, we employed a lipidomic approach. Lipids were extracted from the BAL supernatant and cell pellet of CD44^+/+^ and CD44^-/-^ mice and subjected to LC-MS analysis. From principal component analysis, the lipids from the cell pellet showed a greater difference between the CD44^+/+^ and CD44^-/-^ samples, than the lipids from the BAL supernatant (Figure 3A-B). The volcano plots showed a general trend of more lipids present in the CD44^-/-^ samples, with greater differences observed in the cell pellet (Figure 3C-D). There were 429 lipid moieties identified from the BAL supernatant, and out of these, 52 were significantly changed (p <0.05) by 1.5-fold or more in the CD44^-/-^ samples. The majority of these, 49/52, were increased in the CD44^-/-^ AMs, typically by 1.5 to 2-fold (Supplemental Table 2). The five most significantly changed lipids were 3 moieties of PC and lysophosphatidylglycerol (LPG) which were upregulated, while phosphatidyl serine (PS) was downregulated 4-fold. In the BAL cell pellet, 658 lipids moieties were detected, and out of these, 241 were significantly different: 186 lipids were more abundant and 55 were less abundant in the CD44^-/-^ samples (Supplemental Table 3). In these samples the fold differences varied from 10-fold less to 18 times more. Six out of the top ten most significant changes were increases in PC moieties in the CD44^-/-^ cell pellet. Over five-fold increases were observed with specific TAG, DAG, PA, PC, PG, and PE moieties. Notably, specific oxidized phospholipids (OxPLs) were increased up to 11-fold in the CD44^-/-^ cell pellet, and cholesterol was 2.6-fold higher. Overall, there were many lipid moieties that were present at higher levels in the CD44^-/-^ BAL supernatant and cell pellet. When the lipids were grouped into their main classes and compared, the overall differences in the BAL supernatant were small, with a 1.1 to 1.5-fold increase in the CD44^-/-^ samples, yet significant changes occurred for PC and lysophospholipids. Larger increases (1.7 to 2.6-fold) were observed in the CD44^-/-^ cell pellet for cholesterol and the most prevalent lipids: PC, PE, PG and fatty acids, with 3.6-fold greater levels observed for OxPLs (Figure 3E).

**Figure 3.**
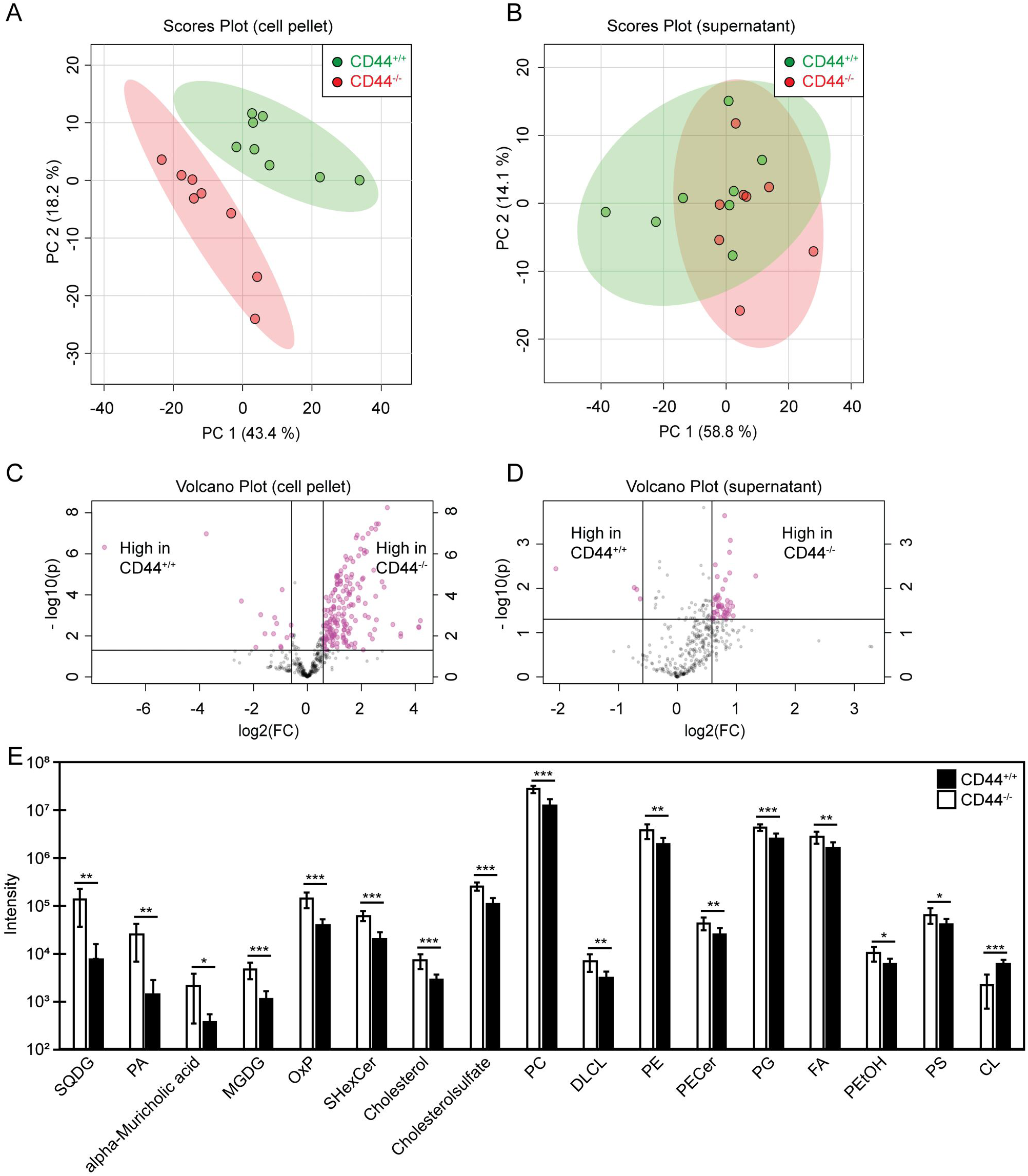
Univariant and multivariant analysis of the differences in lipid composition between CD44^+/+^ and CD44^-/-^ BAL samples. **(A)** and **(B)** Principal component analysis showing lipid differences from the BAL cell pellets and supernatants from CD44^+/+^ and CD44^-/-^ mice. **(C)** and **(D)** Volcano plots comparing the differences in lipid levels measured from the BAL cell pellets and supernatants from CD44^+/+^ and CD44^-/-^ mice. Pink dots represent lipid species with greater than 1.5-fold difference between CD44^+/+^ and CD44^-/-^ samples, with p-value < 0.05. **(E)** Graph showing lipids and lipid classes that are significantly different between CD44^+/+^ and CD44^-/-^ BAL cell pellets. Data show the average from 8 CD44^+/+^ and 8 CD44^-/-^ female mice. Significance indicated as * p< 0.05, ** p< 0.01, *** p < 0.001, using a two tails distribution, homoscedastic *t* test in Microsoft Excel.

### CD44^-/-^ AMs have more lipid droplets, increased ROS and higher OxPC levels

To corroborate the findings from the lipidomic experiments, PC and cholesterol levels were measured by ELISA and gas chromatography respectively, and both were significantly increased in the BAL supernatant from CD44^-/-^ mice (Figure 4A-B). In addition, BODIPY 493/503, a dye that labels neutral lipids/lipid droplets was used to measure lipid levels in AMs. By confocal microscopy, we observed an increase in the number and size of lipid droplets in CD44^-/-^ AMs, and by flow cytometry we observed a significant increase in MFI of BODIPY, indicating a higher neutral lipid content (Figure 4C-D). These data further support a buildup of lipids in the BAL supernatant and AMs from CD44^-/-^ mice.

**Figure 4.**
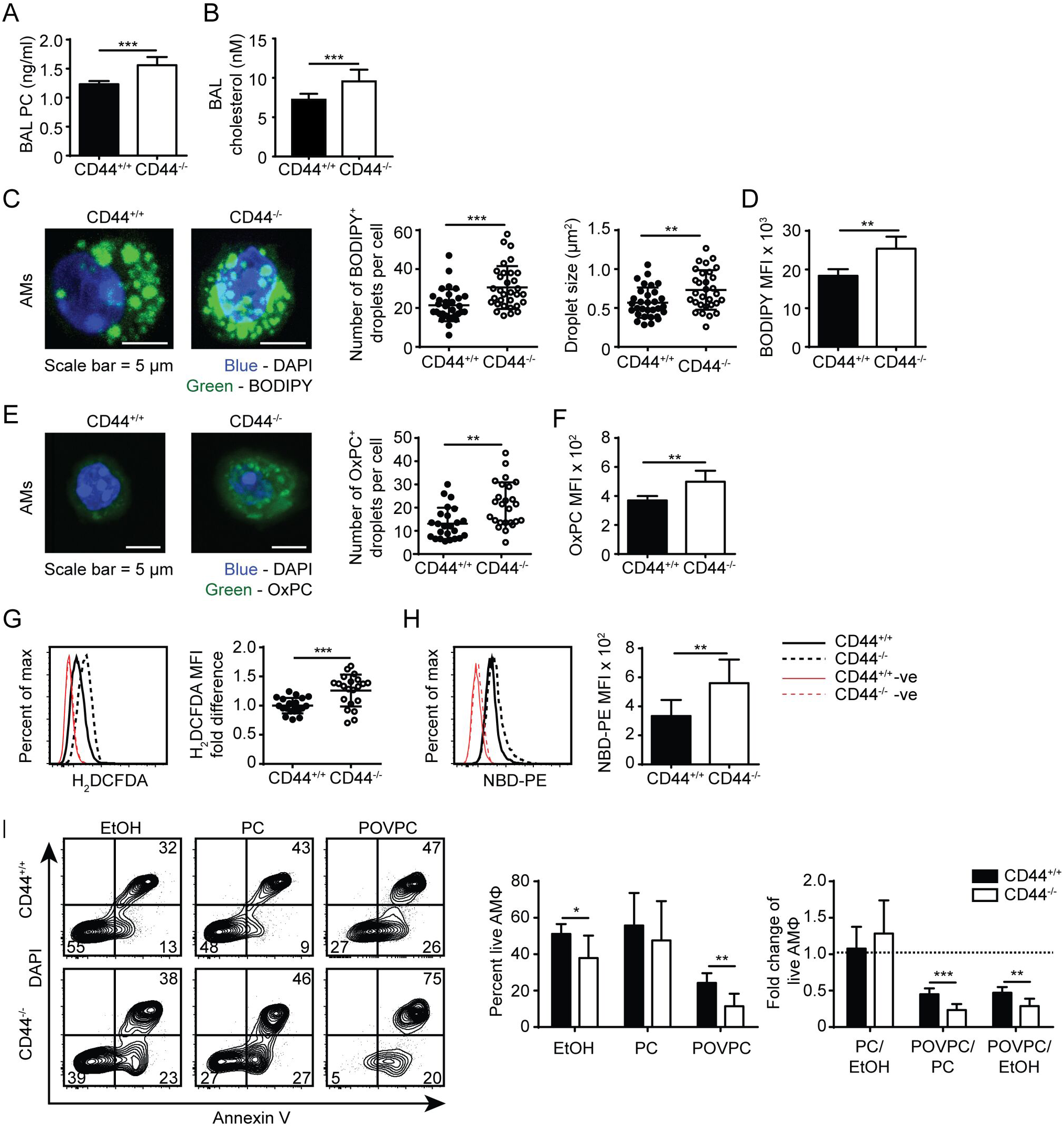
Lipid droplet and oxidized PC levels are elevated in CD44^-/-^ AMs. **(A)** Surfactant PC levels in the BAL supernatant of CD44^+/+^ and CD44^-/-^ mice measured by ELISA. **(B)** Concentration of cholesterol in BAL from CD44^+/+^ and CD44^-/-^ mice measured by thin layer chromatography. **(C)** Representative confocal microscopy (z-stack) of ex vivo CD44^+/+^ and CD44^-/-^ AMs labeled with BODIPY and DAPI, and the number and size of BODIPY^+^ droplets per CD44^+/+^ or CD44^-/-^ AM. **(D)** MFI of BODIPY labeled CD44^+/+^ and CD44^-/-^ AMs by flow cytometry, after subtraction of autofluorescence MFI. **(E)** Representative confocal microscopy (z-stack) of ex vivo CD44^+/+^ and CD44^-/-^ AMs labeled with the anti-OxPC antibody, E06, and DAPI, and the number of E06^+^ droplets per CD44^+/+^ or CD44^-/-^ AM. **(F)** MFI of OxPC levels in CD44^+/+^ and CD44^-/-^ AMs (after subtraction of autofluorescence MFI) using the anti-OxPC antibody E06 and flow cytometry. **(G)** Representative histograms and graphs comparing H2DCFDA labeling of ROS and **(H)** NBD-PE in CD44+/+ and CD44^-/-^ AMs. **(I)** Representative flow cytometry plots of CD44^+/+^ and CD44^-/-^ AMs treated with EtOH, PC, or POVPC labeled with Annexin V and DAPI. Bar graphs show the percent live CD44^+/+^ or CD44^-/-^ AMs (Annexin V^-^ DAPI^-^) after EtOH, PC, or POVPC treatment for 1 h, and compares the fold change in percent live CD44^+/+^ or CD44^-/-^ AMs with different treatments. Data show an average of three to five CD44^+/+^ and CD44^-/-^ mice from each experiment ± SD, repeated twice, or five times **(G)**; three to four cells were analyzed per mouse in confocal imaging, from three to five mice per experiment, over two experiments. Significance indicated as * p< 0.05, ** p< 0.01, *** p < 0.001, non-paired Student’s t-test.

To further investigate the increased presence of oxidized phospholipids in CD44^-/-^ AMs, we used the E06 antibody that recognizes oxidized phosphatidylcholine (OxPC) (38). By confocal microscopy there was an increased number of OxPC droplets per cell and flow cytometry revealed a greater MFI in the CD44^-/-^ AMs (Figure 4E-F). Labeling the AMs with a cell permeant fluorogenic dye that measures cellular ROS activity (H2DCFDA), showed a small but significant increase in ROS activity in the CD44^-/-^ AMs compared to the CD44^+/+^ AMs(Figure 4G). An increased accumulation of lipids and oxidized lipids in the cell are signs of a foamy macrophage and phospholipidosis (39), and this was further supported by the increased labeling of NBD-PE by CD44^-/-^ AMs (Figure 4H). To determine if the increased lipid and OxPC accumulation in CD44^-/-^ AMs resulted in increased oxidized lipid-induced toxicity, CD44^+/+^ and CD44^-/-^ AMs were incubated with the oxidized phospholipid, POVPC, *in vitro* and compared with PC or the vehicle control (1% ethanol). The loss in AM viability caused by POVPC relative to ethanol or PC treatment was significantly greater in CD44^-/-^ AMs compared to CD44^+/+^ AMs (Figure 4I), indicating increased sensitivity to OxPC-induced cellular toxicity.

### The inflammatory response to OxPC is exacerbated in CD44^-/-^ mice

During an inflammatory response, there is increased oxidation of lung surfactant lipids, which can be cytotoxic to AMs if they accumulate. This oxidized lipid-induced cytotoxicity causes lung damage and further drives the inflammatory response (15, 16). To determine if the increased sensitivity of CD44^-/-^ AMs to OxPC observed *in vitro* translated into a more severe inflammatory response in CD44^-/-^ mice, we used a mouse model of OxPC-driven lung inflammation. POVPC was delivered i.t. to CD44^+/+^ and CD44^-/-^ mice and their weight loss was monitored over time (Figure 5A). Analysis of the cells in the BAL 3 days after POVPC instillation, showed leukocyte infiltration including monocytes, neutrophils, eosinophils, and CD11b^-^ cells, indicative of an inflammatory response in both CD44^+/+^ and CD44^-/-^ mice (Figure 5B). Apart from the tissue resident AMs, there were significantly greater numbers of other leukocytes (neutrophils, monocytes, eosinophils) in the BAL from CD44^-/-^ mice (Figure 5C). Protein and HA levels, both indicators of lung inflammation and damage (23, 28, 29), were also increased after POVPC treatment and were higher in the BAL from CD44^-/-^ mice (Figure 5D-E). There was also increased HA detected by HABP labeling in the lung tissue of CD44^-/-^ mice 3 days after POVPC treatment, a sign of increased lung inflammation (Figure F-G). Thus, the increased sensitivity to OxPC toxicity in CD44^-/-^ AMs leads to increased lung damage and greater lung inflammation.

**Figure 5.**
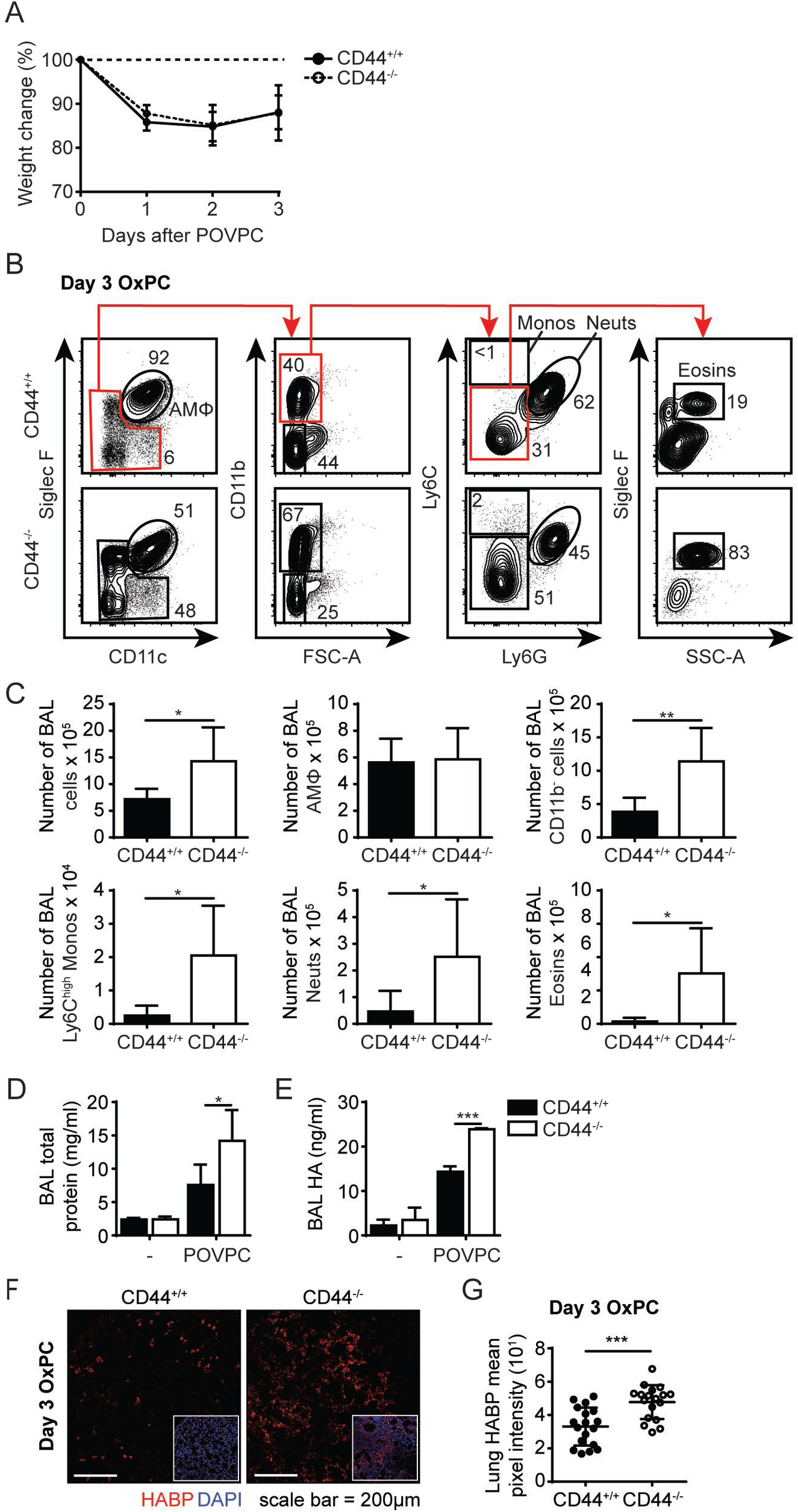
POVPC induced pulmonary inflammation is exacerbated in CD44^-/-^ mice. **(A)** Percent weight change over 3 days from CD44^+/+^ and CD44^-/-^ mice treated with i.t. POVPC. **(B)** Representative flow cytometry plots showing the gating (after live/dead and sizing gating) and proportion of AMs, Ly6C^high^ monocytes, neutrophils, eosinophils, and CD11b^-^ cells in the BAL of CD44^+/+^ and CD44^-/-^ mice 3 days after POVPC instillation. **(C)** Graphs comparing the number of total BAL cells, AMs, CD11b^-^ cells, Ly6c^high^ monocytes, neutrophils, and eosinophils in the BAL of CD44^+/+^ and CD44^-/-^ mice 3 days after POVPC instillation **(D)** Level of total protein in the BAL supernatant of CD44^+/+^ and CD44^-/-^ mice treated with or without POVPC measured by BCA. **(E)** Level of HA in the BAL of CD44^+/+^ and CD44^-/-^ mice treated with or without POVPC measured by ELISA. **(F)** Confocal microscopy and **(G)** graphs comparing HABP mean pixel intensity from CD44^+/+^ and CD44^-/-^ lung sections at 3 days after POVPC treatment. Data show an average of two experiments ± SD, each with three to five CD44^+/+^ and CD44^-/-^ mice. Data from confocal microscopy are pooled from eighteen CD44^+/+^ and CD44^-/-^ POVPC lung sections, from two experiments each with three to five mice. Significance indicated as * p< 0.05, ** p< 0.01, *** p < 0.001, non-paired Student’s t-test and **(G only**) Welch’s t-test with Welch’s correction.

### The effect of the alveolar environment on CD44^-/-^ AMs

To further understand the cause of this lipid accumulation in the AMs and the BAL of CD44^-/-^ mice, we sought to distinguish between intrinsic and extrinsic AM factors. To address the contributions of the alveolar environment on the AMs, we performed short-term adoptive transfer of CD44^-/-^ AMs into the alveolar space of CD44^+/+^ mice and *vice versa*. CD44^-/-^AMs expressing the CD45.2 allele, were instilled into the trachea of CD44^+/+^ BoyJ mice expressing the CD45.1 allele and the BAL analyzed seven days later (Figure 6A). After being transferred to a normal alveolar environment, the CD45.2^+^ CD44^-/-^ AMs decreased expression of CD36 (Figure 6B-C), consistent with CD36 expression responding to the reduced lipid surfactant environment of the CD44^+/+^ mice. However, BODIPY, SSC-A and CD11c levels did not drop significantly. In contrast, when CD45.1^+^ CD44^+/+^ AMs were transferred into the trachea of the CD45.2^+^ CD44^-/-^ mice and analyzed seven days later (Figure 6D), these AMs dramatically increased CD36 expression, to levels higher than the CD44^-/-^ AMs (Figure 6E-F), consistent with increased lipid surfactant levels in the CD44^-/-^ mice. In addition, SSC-A and BODIPY levels increased to similar levels observed in CD44^-/-^ AMs, however, the levels of CD11c did not increase. To evaluate if the CD44^+/+^ AMs were directly responding to increased lipid levels in the lung surfactant of CD44^-/-^ mice, *ex vivo* CD44^+/+^ AMs were incubated with PC for 48 h *in vitro* then analyzed for neutral lipids using BODIPY, and CD36 expression. Both showed a significant increase (Figure 6G-I), demonstrating a direct correlation between extracellular PC, CD36 and BODIPY levels (intracellular lipid accumulation) in CD44^+/+^ AMs. This suggests an intrinsic defect in CD44^-/-^ AMs that prevents the decrease in intracellular lipid levels.

**Figure 6:**
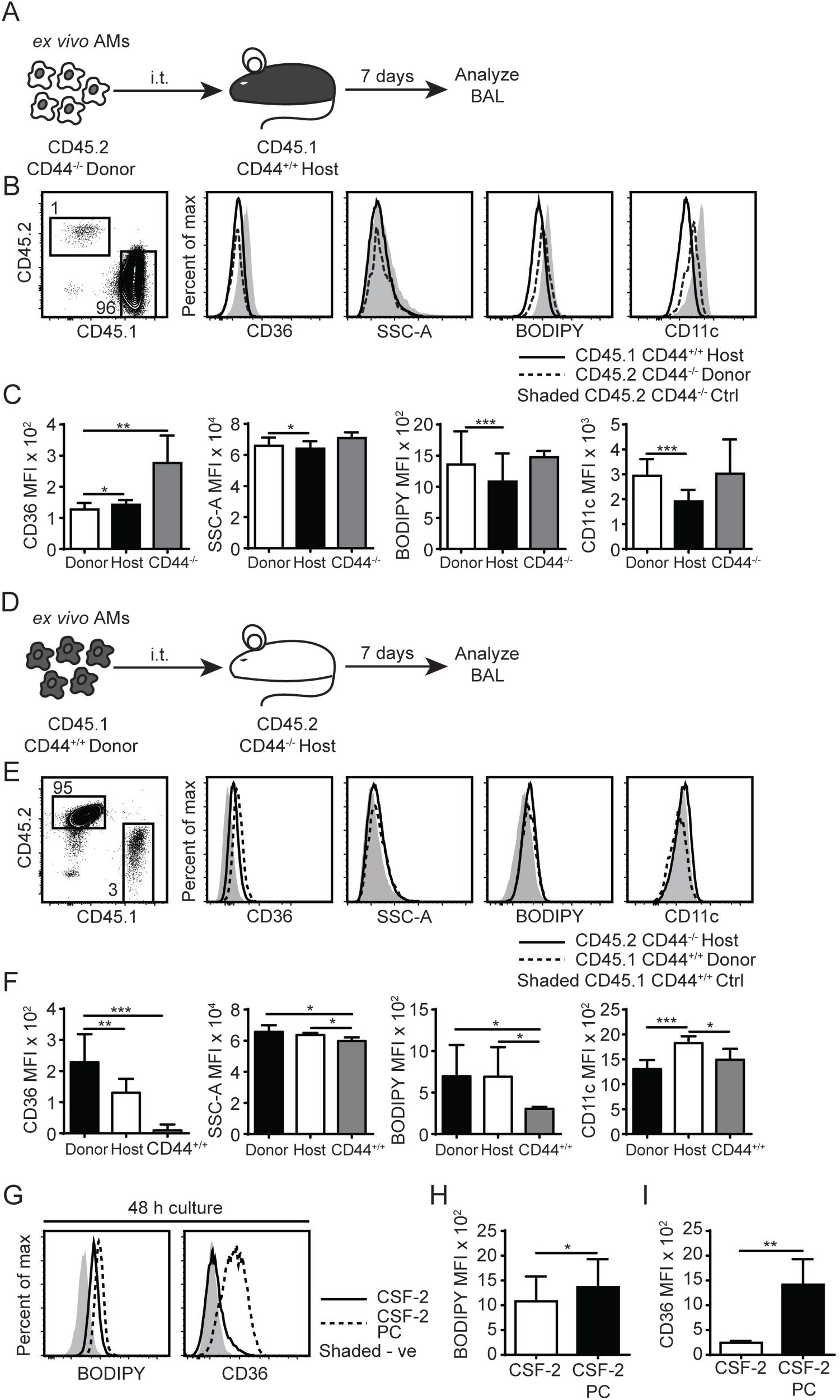
AMs lipid homeostasis is controlled by the extracellular milieu and the expression of CD44. **(A)** Schematic diagram showing donor AMs isolated from CD45.2^+^ CD44^-/-^ mice are transferred by i.t. into CD45.1^+^ CD44^+/+^ host mice and analyzed 7 days later. **(B)** Representative flow cytometry plots comparing the SSC-A, BODIPY, CD36, and CD11c levels by CD45.2^+^ CD44^-/-^ donor AMs and CD45.1^+^ CD44^+/+^ host AMs with control CD45.2^+^ CD44^-/-^ mice, 7 days after adoptive transfer. **(C)** Graphs comparing SSC-A, BODIPY, CD36, and CD11c MFI of CD45.2^+^ CD44^-/-^ donor AMs, CD45.1^+^ CD44^+/+^ host AMs, and ex vivo CD44^-/-^ AMs. **(D)** Schematic diagram showing donor AMs isolated from CD45.1^+^ CD44^+/+^ mice are transferred by i.t. into CD45.2^+^ CD44^-/-^ host mice and analyzed 7 days later. **(E)** Representative flow cytometry plots comparing the SSC-A, BODIPY, CD36, and CD11c levels in CD45.1^+^ CD44^+/+^ donor AMs and CD45.2^+^ CD44^-/-^ host AMs compared with CD45.1^+^ CD44^+/+^ control mice, 7 days after adoptive transfer. **(F)** Graphs comparing SSC-A, BODIPY, CD36, and CD11c MFI of CD45.1^+^ CD44^+/+^ donor AMs, CD45.2^+^ CD44^-/-^ host AMs, and ex vivo CD44^+/+^ AMs. **(G)** Representative flow cytometry histograms comparing the BODIPY and CD36 MFI of AMs after 48 h culture in CSF-2, or CSF-2 and PC. **(H)** and **(I)** Graphs comparing BODIPY labeling and CD36 expression by AMs cultured in CSF-2, or CSF-2 and PC for 48 h. For adoptive transfer experiments, data show an average of two experiments ± SD, each with three to five mice. Significance indicated as * p< 0.05, ** p< 0.01, *** p < 0.001, paired (donor AMs vs host AMs) and non-paired Student’s t-test (CD44^-/-^ donor AMs vs ex vivo CD44^-/-^ AMs or CD44^-/-^ host AMs vs ex vivo CD44^+/+^ AMs). For in vitro cell culture experiments, data show an average of two experiments ± SD, each with three to five mice. Significance indicated as * p< 0.05, ** p< 0.01, *** p < 0.001, paired Student’s t-test.

To further demonstrate cell intrinsic defects, we performed bone marrow reconstitution experiments on irradiated mice. We reconstituted irradiated CD45.1^+^ CD44^+/+^ mice with CD45.2^+^ CD44^-/-^ bone marrow (Figure 7A). After allowing several weeks for reconstitution of the AM compartment, CD44^-/-^ AMs were isolated from the BAL and had a similar phenotype to CD44^-/-^ AMs isolated from CD44^-/-^ mice, namely similar levels of BODIPY, CD36 and CD11c (Figure 7B-C). We also reconstituted irradiated CD44^+/+^ mice with 50% CD44^+/+^ and 50% CD44^-/-^ bone marrow and analyzed both CD44^+/+^ and CD44^-/-^ AMs from the same lung environment after allowing at least 7 weeks for reconstitution of the lung macrophage compartment (Figure 7D). This competitive transfer leads to the dominance of CD44^+/+^ AMs over CD44^-/-^ AMs in an 80:20 ratio, as CD44^-/-^ AMs are less able to survive (12). Analysis of both the CD44^+/+^ and CD44^-/-^ AMs from the same mice showed that the CD44^-/-^ AMs were significantly higher in SSC and CD11c levels (Figure 7E). Together, these results show the ability of CD44^-/-^ AMs to regulate CD36 expression in response to changes in the lung environment, but an inability to decrease their lipid content or reduce CD11c levels, indicating both cell extrinsic and intrinsic changes in CD44^-/-^ AMs.

**Figure 7.**
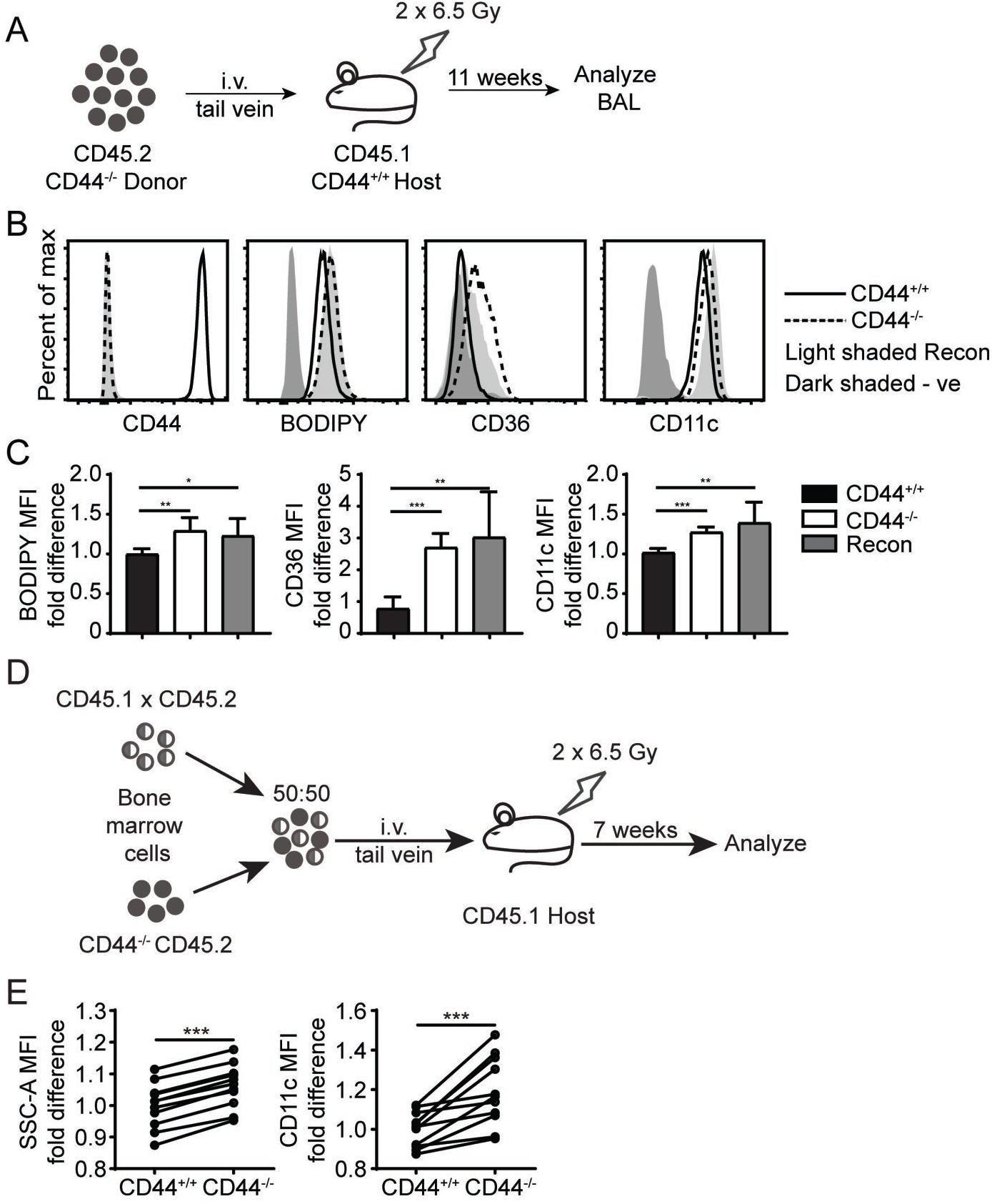
Cell intrinsic defects in CD44^-/-^ AMs are preserved after long term BM reconstitution. **(A)** Schematic showing the BM reconstitution experiment where BM cells from CD44^-/-^ mice were injected into lethally irradiated CD44^+/+^ host mice and analyzed 11 weeks later. **(B)** Representative flow cytometry plots comparing the CD44, BODIPY, CD36, and CD11c levels of AMs from CD44^+/+^ mice and CD44^-/-^ mice, as well CD44^-/-^ AMs reconstituted in CD44^+/+^ mice. **(C)** Graphs comparing the fold difference of BODIPY, CD36, and CD11c of AMs from CD44^+/+^ mice and CD44^-/-^ mice, as well CD44^-/-^ AMs reconstituted in CD44^+/+^ mice. **(D)** Schematic showing the BM reconstitution experiment where BM cells from CD44^+/+^ and CD44^-/-^ mice were injected at equal proportions into lethally irradiated host mice to study the role of CD44 in AM repopulation. **(E)** Graphs comparing the fold difference in the granularity (SSC-A) and CD11c between CD44^+/+^ and CD44^-/-^ AMs in the BAL after 7 weeks in competition, as measured by flow cytometry. Data show an average of two experiments ± SD, each with three to five mice. Significance indicated as * p< 0.05, ** p< 0.01, *** p < 0.001 unpaired Student’s t-test.

### CSF-2 partially restored lipid surfactant homeostasis in CD44^-/-^ BAL but did not reduce lipid droplets in CD44^-/-^ AMs

While the reduced numbers of CD44^-/-^ AMs (12) could contribute to increased PC and cholesterol levels in the BAL, the adoptive transfer and competitive reconstitution experiments suggested an intrinsic defect in the ability to catabolize lipids in the CD44^-/-^ AMs, which was also supported by the lipidomic data. Lipid droplet accumulation and the foamy AM phenotype is a feature of immature AMs that arise after the deletion of CSF-2, PPARγ or TGFβ, which are important for maturation and survival (2-5). While the RNAseq data showed reduced TGFβ2 transcript levels in CD44^-/-^ AMs, these transcripts were very low compared to the levels of TGFβ1 transcripts, which did not differ between normal and CD44^-/-^ AMs (geo accession no: GSE138445). CSF-2 induces PPARγ expression (4) and upregulates CD11c expression in bone marrow derived macrophages (12), and both CSF-2 and PPARγ agonists promote CD44-mediated HA binding (12). It is possible that some of the downstream effects of CSF-2 or PPARγ could be mediated via CD44 and its interaction with HA, which then promotes AM survival (12). Since CSF-2 also attenuates bleomycin-induced lipid accumulation in the alveolar space (16), we determined if CSF-2 could rescue the defects in surfactant lipid homeostasis caused by CD44 deficiency. CSF-2 was given daily to the CD44^-/-^ mice by i.t. for seven days, then the BAL from the mice was analyzed. While CSF-2 instillation did not significantly alter the composition of cells in the BAL (Figure 8A), it did increase CD44^-/-^ AM numbers (Figure 8B) and decrease the concentration of PC in the BAL (Figure 8C). There was a concomitant reduction in CD36 expression in the CD44^-/-^ AMs, but no significant changes were observed in their granularity (SSC-A), neutral lipid content (measured by BODIPY), or the expression of CD11c (Figure 8D-E). This suggests that the reduced number of AMs contributes to increased lipid surfactant levels and concomitantly, increased levels of CD36. However, CSF-2 does not rescue the cell intrinsic defect of accumulated lipids in CD44^-/-^ AMs.

**Figure 8.**
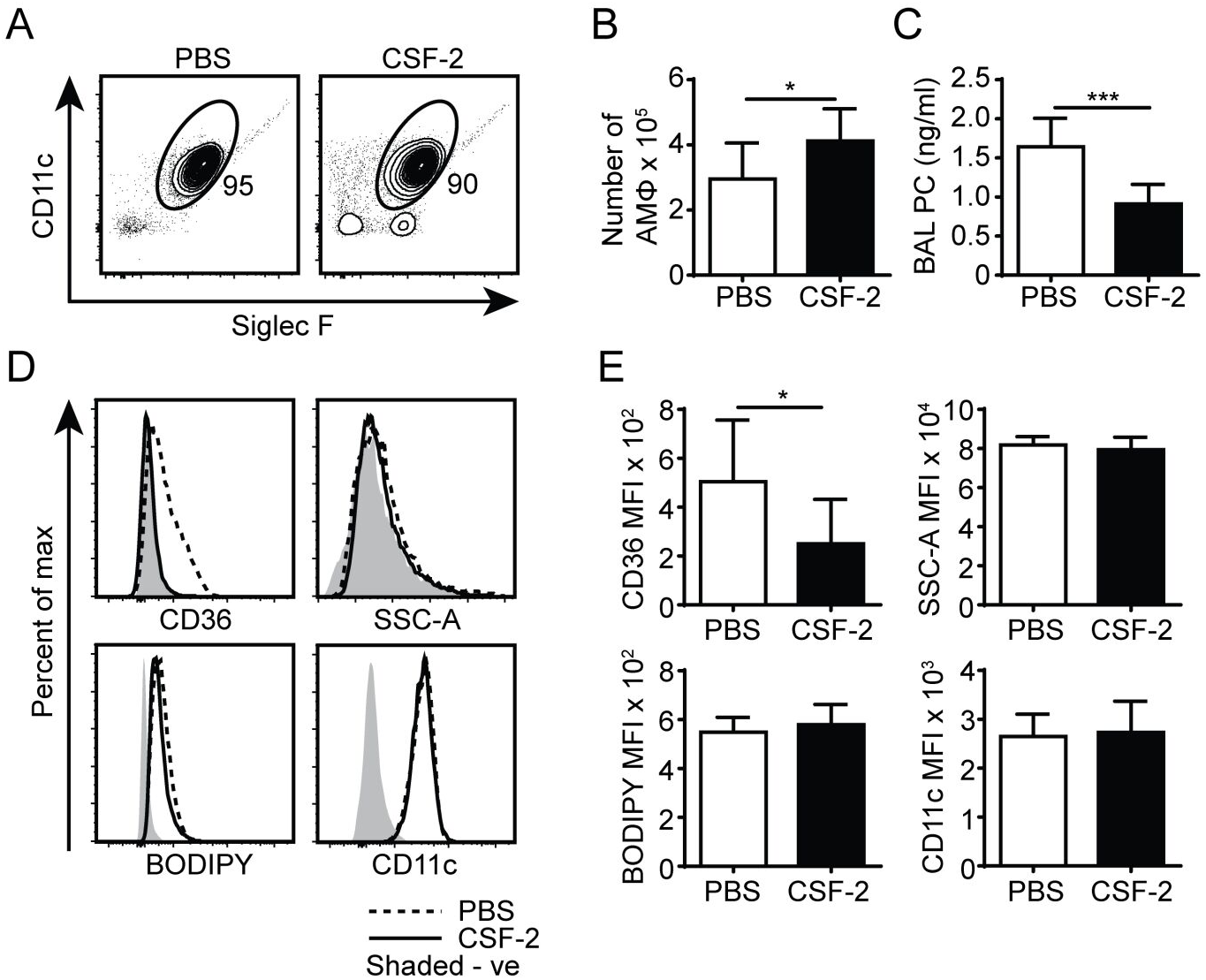
CSF-2 induces AM proliferation and rescues aberrant pulmonary surfactant PC accumulation in CD44^-/-^ mice. **(A)** Representative flow cytometry plots showing the proportion of BAL AMs from CD44^-/-^ mice after 7 days of PBS or CSF-2 treatment. **(B)** Number of AMs in the BAL from CD44^-/-^ mice after PBS or CSF-2 treatment. **(C)** BAL PC concentration from CD44^-/-^ mice after PBS or CSF-2 treatment. **(D)** Representative flow cytometry histograms comparing the phenotype of AMs from CD44^-/-^ mice after 7 days of PBS or CSF-2 treatment. **(E)** Graphs comparing the MFI of SSC-A, BODIPY, CD36, and CD11c between AMs from CD44^-/-^ mice after 7 days of PBS or CSF-2 treatment. Data show an average of two experiments ± SD, each with three to five mice. Background MFI of autofluorescence from unlabeled CD44^+/+^ and CD44^-/-^ cells were respectively subtracted in the flow cytometry analysis. Significance indicated as * p< 0.05, *** p < 0.001, non-paired Student’s t-test.

### CD44^-/-^ AMs have reduced PPARγ expression

PPARγ is an important transcriptional regulator of lipid metabolism (10) and this, coupled with its key role in AM maturation and survival (4), make it a potential candidate for contributing to the cell intrinsic defects present in the CD44^-/-^ AMs. Although no differences in PPARγ transcript levels were apparent from the RNAseq data, other forms of regulation such as PPARγ activation, cellular localization, or protein levels may be defective in CD44^-/-^ AMs. Confocal microscopy showed that the majority of PPARγ in *ex vivo* CD44^+/+^ and CD44^-/-^ AMs was localized to the nucleus (Figure 9A). Both confocal microscopy and flow cytometry showed PPARγ protein was expressed at a slightly lower level in CD44^-/-^ AMs (Figure 9B-C). To determine if this was of functional significance, *ex vivo* CD44^+/+^ AMs were incubated for 48 h in the presence of CSF-2 and treated with a selective PPARγ antagonist, T-0070907 (40), to mimic conditions in the CD44^-/-^ AMs. The antagonist increased the levels of BODIPY, as well as CD36 and CD11c (Figure 9D-E). This suggests that the reduced expression/activation of PPARγ could contribute to the accumulation of lipids in CD44^-/-^ AMs.

**Figure 9.**
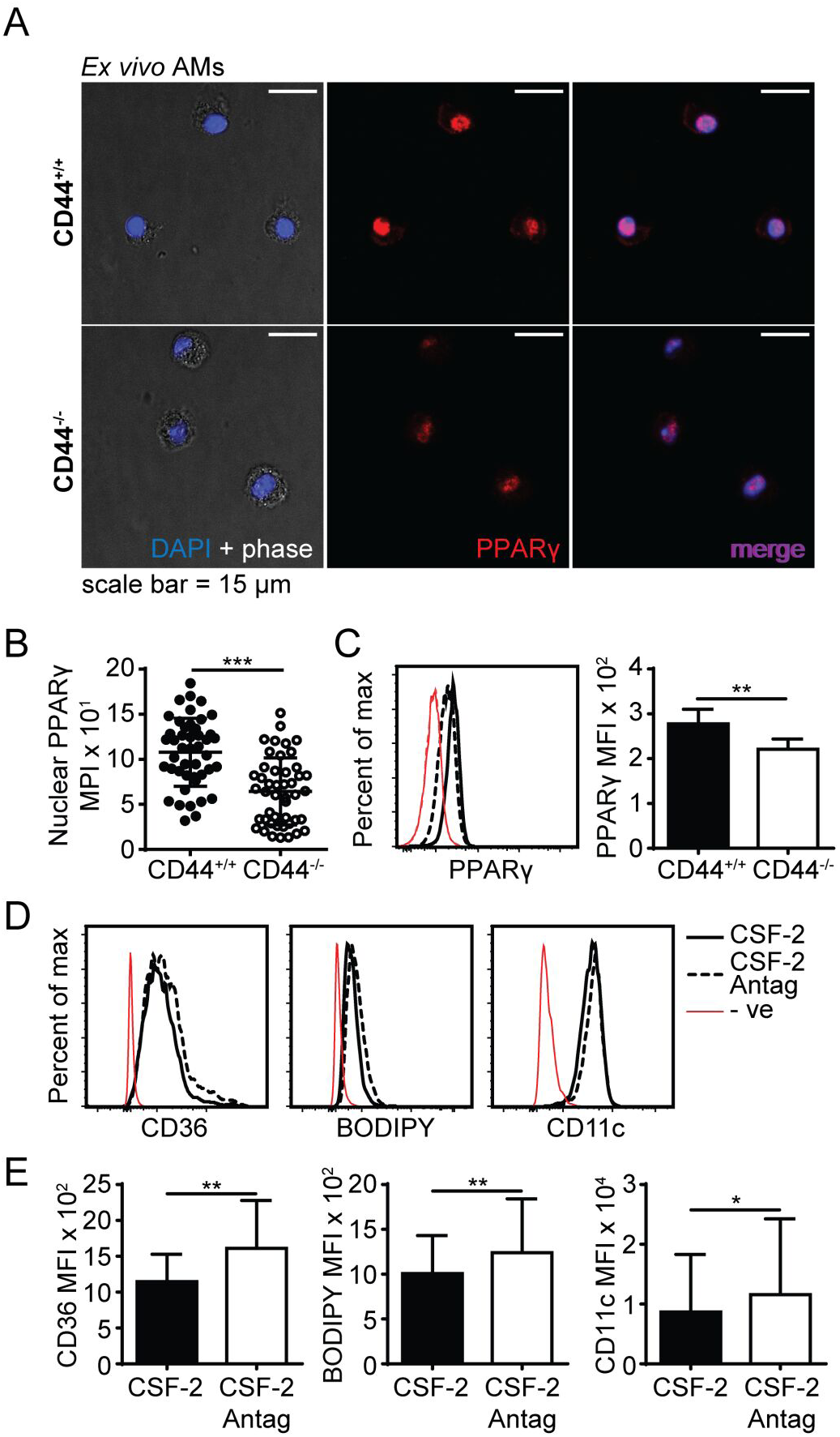
PPARγ expression is defective in CD44^-/-^ AMs. **(A)** Representative confocal microscopy showing CD44^+/+^ and CD44^-/-^ AMs labeled with intracellular PPARγ antibody and DAPI. **(B)** Comparison of nuclear PPARγ mean pixel intensity (MPI) between CD44^+/+^ and CD44^-/-^ AMs, determined by confocal microscopy. **(C)** Representative flow cytometry histograms and graphs comparing intracellular PPARγ expression between CD44^+/+^ and CD44^-/-^ AMs. **(D)** and **(E)** Representative flow cytometry histograms and graphs comparing the levels of CD36, BODIPY and CD11c levels by MFI (after subtraction of background autofluorescence) in CD44^+/+^ AMs cultured with CSF-2 or CSF-2 and the PPARγ antagonist T0070907 (Antag) for 48 h in culture. Data show an average of three to five mice from each experiment ± SD, repeated twice or three times; for confocal imaging three to four fields containing cells were analyzed per mice. Significance indicated as * p< 0.05, ** p< 0.01, *** p < 0.001, non-paired Student’s t-test.

## DISCUSSION

AMs play an important role in maintaining lung homeostasis and are responsible for the uptake of surfactant and oxidized surfactant lipids, their storage and catabolism (1). Here we report that CD44 deficiency disrupted lipid surfactant homeostasis in the alveolar space and led to intracellular lipid accumulation in AMs. CD44^-/-^ AMs accumulated PC and cholesterol and had increased intracellular lipid droplets, giving the AMs a foamy appearance, with increased side scatter (SSC) and autofluorescence. CD44 deficiency in AMs led to the differential expression of 200 genes, several of which were involved in lipid metabolism and trafficking. The inability of CD44^-/-^ AMs to reduce intracellular lipid levels coupled with the reduced numbers of AMs in CD44^-/-^ mice resulted in the increase in lung surfactant lipids in the BAL supernatant. The inability to reduce intracellular lipid accumulation in CD44^-/-^ AMs may be attributed, at least in part, to the dysregulation of PPARγ.

Transcriptional analysis of CD44^+/+^ and CD44^-/-^ AMs identified the upregulation of genes involved in cholesterol efflux and trafficking in CD44^-/-^ AMs. These included *ApoE* (41), which was validated by qPCR (data not shown) and *ABCA1* (42), key genes downstream of PPARγ signaling. Although *CD36* gene levels were not significantly different by transcriptomic analysis, cell surface expression of CD36 was increased in CD44^-/-^ AMs, and this correlated with extracellular lipid levels. CD36 is a multi-functional scavenger receptor that binds oxidized lipids and fatty acids (37) and CD36 signaling promotes cholesterol efflux and inhibits cholesterol synthesis by activating PPARγ (43). The transcriptional analysis also predicted the down regulation of several genes in CD44^-/-^ AMs, including *srebf2*, a master transcriptional regulator of cholesterol and fatty acid synthesis (44, 45) and a cluster of genes associated with sterol/cholesterol synthesis. Together, these data suggest that CD44^-/-^ AMs are responding to increased exposure to lipids by activating gene pathways responsible for reducing lipid content, yet are still unable to reduce intracellular lipid content. Although PPARγ is a master transcriptional regulator of lipid metabolism (8-10), its effects can be cell-type specific and complicated by the activation of its heterodimeric partner, RXR and other lipid regulated transcription factors such as LXR. While one may have predicted that PPARγ would have been more active in CD44^-/-^ AMs, transcript levels were not different and slightly less, not more, PPARγ protein levels were found in the nucleus of CD44^-/-^ AMs.

In AMs, PPARγ expression is induced by CSF-2 and TGFβ and is critical for their development (4). PPARγ upregulates genes important for the transcriptional identity of AMs, including molecules involved in the intracellular binding, storage and metabolism of lipids, which help prevent foam-cell formation and protect against lipotoxicity (4, 9, 10, 46). PPARγ deficiency in AMs causes significant accumulation of cholesterol and phospholipids in the BAL supernatant (47, 48) and leads to immature AMs that have increased lipid droplets (4). This is consistent with the results here, where we found slightly reduced levels of PPARγ in the nucleus of CD44^-/-^ AMs. The upregulation of BODIPY^+^ lipid droplets and CD36 on CD44^+/+^ AMs after *in vitro* culture with a PPARγ antagonist also supports the idea that reduced PPARγ activity mimics the CD44^-/-^ AM phenotype. However, in addition to lipid induced activation of PPARγ, its activity can also be regulated at the translational and protein level. PPARγ protein is degraded via ubiquitination after ligand mediated activation (49), and so it is possible that CD44^-/-^ AMs have lower nuclear PPARγ levels because they have been exposed to greater levels of activating PPARγ ligands, leading to greater degradation.

Whether CD44 directly or indirectly modulates PPARγ activity remains to be addressed. In IL-1β stimulated chondrosarcoma cells, the addition of high molecular weight HA increases PPARγ expression at both the mRNA and protein level (50). It will be of interest to determine if there is a link between HA engagement by CD44 and PPARγ levels or activation. Alternatively, TGFβ can upregulate PPARγ expression (4) and RNAseq data did predict a reduction in TGFβ2 expression in CD44^-/-^ AMs, although not for the more highly expressed TGFβ1. In other cell types, CD44 and HA-binding modulate TGFβ responses by facilitating TGFβ cleavage and activation (51), or by promoting TGFβ receptor signaling (52).

Mice deficient in CSF-2, or harboring an AM-specific deletion of PPARγ, or the conditional deletion of TGFβR2, all produce CD11b^+^ immature foamy macrophages, which are unable to regulate lipid turnover leading to surfactant accumulation in the BAL and ultimately PAP (3-5). Their immature CD11b^+^ CD11c^low^ Siglec F^low^ phenotype is distinct from the phenotype of CD44^-/-^ AMs, which have high levels of mature AM markers: Siglec F, CD11c, CD206, CD200R, and Sirpα. However, CD44^-/-^ AMs also express low levels of CD11b and MHCII, and RNAseq analysis predicted an upregulation of *mafB*, a transcription factor that is normally down-regulated in mature cells, to allow self-renewal and terminal differentiation (53). It is possible some of the effects of CSF-2 on AMs maybe mediated by CD44 as CSF-2 induces HA binding by CD44 and this is known to promote the survival of AMs (12). Although exogenous CSF-2 restored AM numbers and reduced PC levels in the BAL of CD44^-/-^ mice, it did not impact the intracellular lipid accumulation in CD44^-/-^ AMs.

RNAseq predicted the altered expression of genes involved in cholesterol/sterol synthesis and trafficking, triglyceride synthesis and phospholipid catabolism in CD44^-/-^ AMs. This suggested dysregulation of lung lipid surfactant metabolism in CD44^-/-^ AMs, which was validated by measuring the lipids present in the BAL supernatant and cell pellet of the CD44^+/+^ and CD44^-/-^ mice. Although the molecular composition of lung surfactant has been analyzed in young and old mice (54), to our knowledge, this study is the first to provide a full lipidomic analysis of the lung surfactant and AMs. The major lipids present in lung surfactant include PC moieties (1, 7, 11), which were elevated in the CD44^-/-^ BAL supernatant and cell pellet. Lipidomic MS analysis of the cell pellets from CD44^+/+^ and CD44^-/-^ AMs showed greater accumulation of intracellular lipids including PC, PE, PG, fatty acids, cholesterol, and OxPLs in CD44^-/-^ AMs. These results were consistent with the observations from flow cytometry and confocal microscopy experiments which showed CD44^-/-^ AMs had higher BODIPY labeled neutral lipids and OxPC levels than CD44^+/+^ AMs. The increased cytotoxicity of OxPC in CD44^-/-^ AMs coupled with the observed increase in ROS and OxPC levels in these cells, resulted in an increased inflammatory response in the lungs of CD44^-/-^ mice exposed to oxidized lipids. This is of significance as oxidized lipids propagate an inflammatory response. Interestingly, mice exposed to bleomycin generate oxidized lipids (16) and CD44^-/-^ mice have significantly worse bleomycin-induced inflammation and lung damage compared to wild-type mice (28).

Overall, this work demonstrates a new role for CD44 in AMs in maintaining lipid surfactant homeostasis, a key function of AMs in the alveolar space. Dysregulation of lipid surfactant homeostasis in CD44^-/-^ AMs leads to a buildup of lipid surfactant and foamy AMs, and an increase in oxidized lipids exacerbates lung damage and inflammation. This work also raises the possibility that reduced numbers of AMs and compromised lipid surfactant metabolism, leading to increased lung damage and inflammation, are contributing factors to the development of secondary pulmonary alveolar proteinosis.

## ACKNOWLEDGEMENTS

We acknowledge invaluable assistance from the UBC Animal and Flow cytometry Facilities.

## AUTHOR CONTRIBUTIONS

YD initiated and developed the project and performed most of the experiments. AA, JH, JG, GP and SL, provided input and performed experiments, and MD provided technical support. CR provided supervision and TH oversaw the lipidomic experiments, provided financial support and supervision. PJ developed and oversaw the project, provided supervision, and acquired funds. YD and PJ wrote the manuscript, with contributions from JG, which was reviewed by all authors.

