## Supplemental figures for "RNA sequencing and lipidomic analysis of alveolar macrophages from normal and CD44 deficient mice"

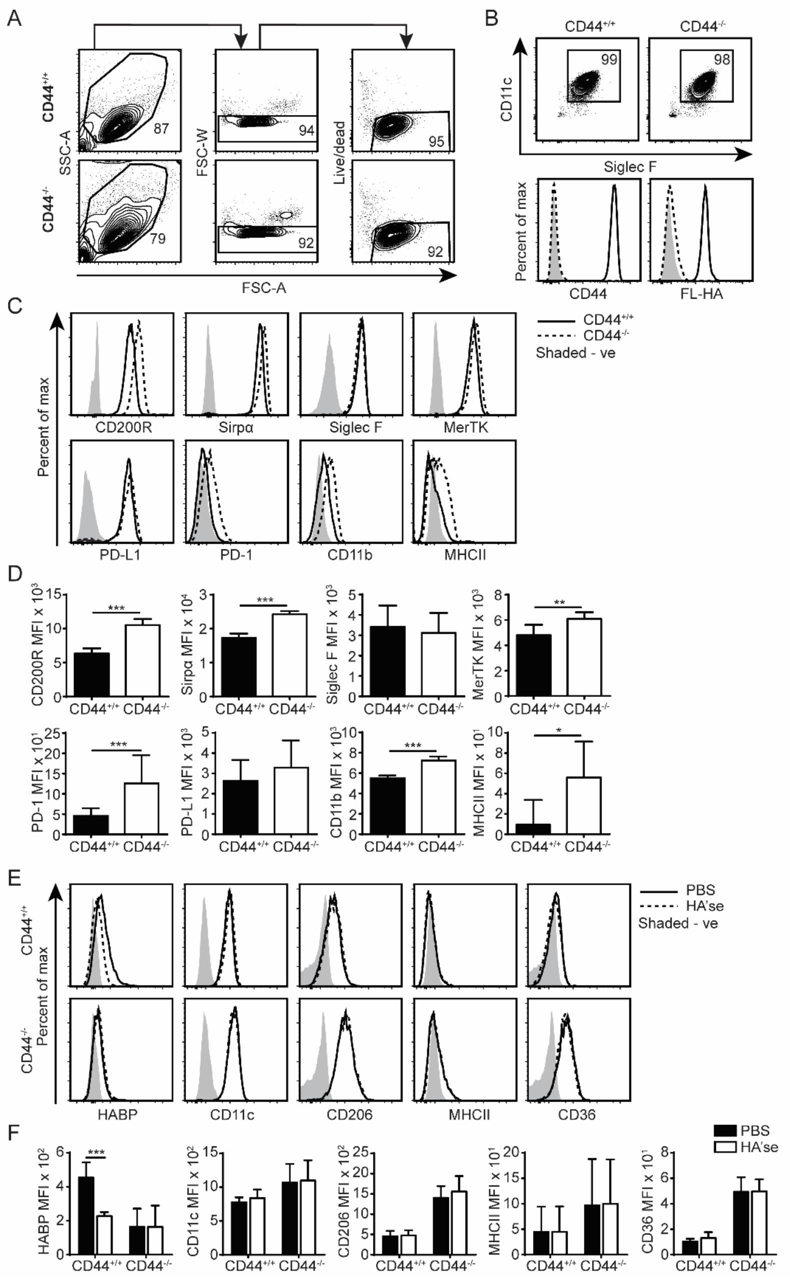


**Supplementary Figure 1.** Comparing the phenotype of CD44$^{+/+}$ and CD44$^{-/-}$ AMs in the presence or absence of cell associated HA. **(A)** and **(B)** Flow cytometry plots showing the gating strategy for cell size, singlets, and live dead cells to identify CD11c$^{+}$ Siglec F$^{+}$ HA binding CD44$^{+/+}$ AMs and HA non-binding CD44$^{-/-}$ AMs from the BAL of CD44$^{+/+}$ and CD44$^{-/-}$ mice, respectively. **(C)** Histogram flow cytometry plots and **(D)** bar graphs comparing the cell surface expression of CD200R, Sirpα, Siglec F, MerTK, PD-1, PD-L1, CD11b, and MHCII between CD44$^{+/+}$ and CD44$^{-/-}$ AMs. **(E)** Histogram flow cytometry plots and **(F)** bar graphs comparing the detection of cell surface hyaluronan binding protein (HABP), CD11c, CD206, MHCII, and CD36 on CD44$^{+/+}$ and CD44$^{-/-}$ AMs with or without the treatment of hyaluronidase (HA'se) to remove cell associated HA. Data show an average of two experiments ± SD, each with three to five CD44$^{+/+}$ and CD44$^{-/-}$ mice. Significance indicated as * $p< 0.05$, ** $p< 0.01$, *** $p < 0.001$, paired (F) and non-paired (D) student's t-test.

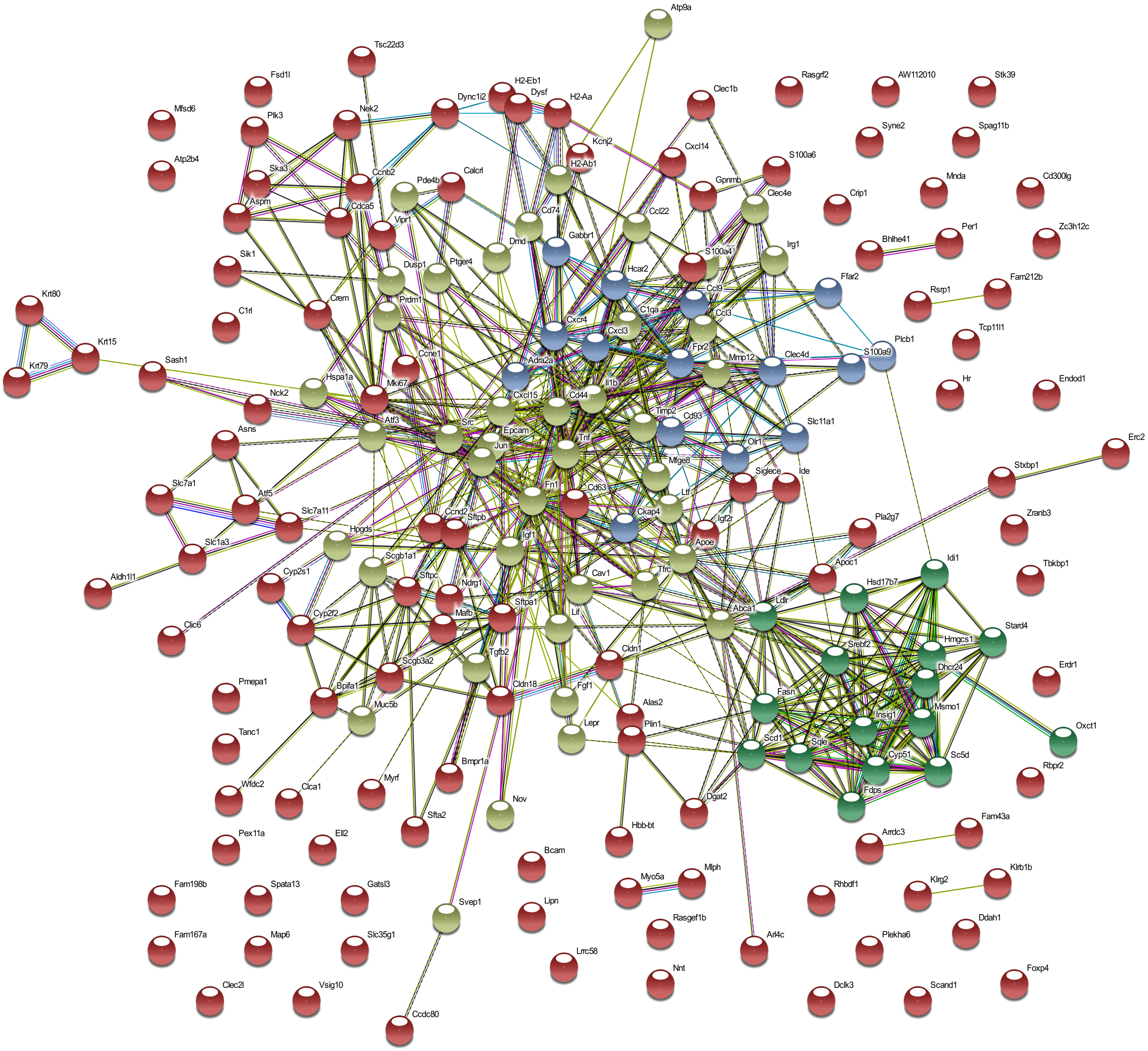


**Supplementary Figure 2.** String analysis of the differentially expressed genes from CD44$^{+/+}$ and CD44$^{-/-}$ AMs acquired by RNA sequencing using the String Consortium (33). Each node represents one of the 200 significant genes (q < 0.05) from Cuffdiff RNA sequencing and is colored according to network clustering which was set to 3. Edges (lines) represent known interactions from curated databases (cyan) or experimentally determined (violet); or predicted associations from gene neighborhood (green), gene fusion (red), and gene co-occurrence (blue); as well as other associations from text mining (olive), co-expression (black), and protein homology (purple).

**Reference (33):** Szklarczyk D, Gable AL, Lyon D, Junge A, Wyder S, Huerta-Cepas J, et al. STRING v11: protein-protein association networks with increased coverage, supporting functional discovery in genome-wide experimental datasets. *Nucleic Acids Res* (2019) 47(D1):D607-D13. doi: 10.1093/nar/gky1131.
